# Pressure sensitivity of ANME-3 predominant anaerobic methane oxidizing community from coastal marine Lake Grevelingen sediment

**DOI:** 10.1101/307082

**Authors:** C. Cassarini, Y. Zhang, P. N. Lens

## Abstract

Anaerobic oxidation of methane (AOM) coupled to sulfate reduction is mediated by, respectively, anaerobic methanotrophic archaea (ANME) and sulfate reducing bacteria (SRB). When a microbial community from coastal marine Lake Grevelingen sediment, containing ANME-3 as the most abundant type of ANME, was incubated under a pressure gradient (0.1-40 MPa) for 77 days, ANME-3 was more pressure sensitive than the SRB. ANME-3 activity was higher at lower (0.1, 0.45 MPa) over higher (10, 20 and 40 MPa) CH_4_ total pressures. Moreover, the sulfur metabolism was shifted upon changing the incubation pressure: only at 0.1 MPa elemental sulfur was detected in a considerable amount and SRB of the <u>*Desulfobacterales*</u> order were more enriched at elevated pressures than the *Desulfubulbaceae*. This study provides evidence that ANME-3 can be constrained at shallow environments, despite the scarce bioavailable energy, because of its pressure sensitivity. Besides, the association between ANME-3 and SRB can be steered by changing solely the incubation pressure.

**Importance:** Anaerobic oxidation of methane (AOM) coupled to sulfate reduction is a biological process largely occurring in marine sediments, which contributes to the removal of almost 90% of sedimentary methane, thereby controlling methane emission to the atmosphere. AOM is mediated by slow growing archaea, anaerobic methanotrophs (ANME) and sulfate reducing bacteria. The enrichment of these microorganisms has been challenging, especially considering the low solubility of methane at ambient temperature and pressure. Previous studies showed strong positive correlations between the growth of ANME and the methane pressure, since the higher the pressure the more methane is dissolved. In this research, a shallow marine sediment was incubated under methane pressure gradients. The investigated effect of pressure on the AOM-SR activity, the formation sulfur intermediates and the microbial community structure is important to understand the pressure influence on the processes and the activity of the microorganisms involved to further understand their metabolism and physiology.

## Introduction

Anaerobic oxidation of methane (AOM) coupled to sulfate reduction (SR) is a major sink in the oceanic methane (CH_4_) budget. The net stoichiometry of this reaction is shown in Eq. 1 (1):

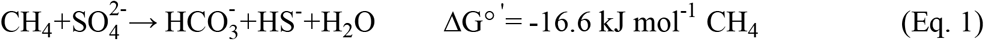

The thermodynamics of this reaction depend on the concentration of dissolved CH_4_. CH_4_ is poorly soluble: 1.3 mM is its concentration in sea water at ambient pressure and at 15°C (2). Theoretically, an elevated CH_4_ pressure favors the AOM coupled to SR (AOM-SR) bioconversion since i) the Gibbs free energy becomes more negative at higher CH_4_ partial pressures and ii) the dissolved CH_4_ concentration increases and is thus more bioavailable (Table 1). Thus, the activity and the growth of the microorganisms mediating the process, namely anaerobic methanotrophs (ANME) and sulfate reducing bacteria (SRB), is expected to be higher at elevated pressures.

ANME are grouped into three distinct clades, i.e. ANME-1, ANME-2 and ANME-3 based on the phylogenetic analysis of their 16S rRNA genes (3–5). *In vitro* incubations of ANME-1 and ANME-2 dominated microbial communities from deep sea sediments in high-pressure reactors showed indeed a strong positive relation of the activity of the microorganisms capable of the AOM-SR process with the CH_4_ partial pressure, up to 12 MPa (6–10). In the ANME-2 dominated shallow marine sediment of Eckernförde Bay, the AOM-SR rate increased linearly with the CH_4_ pressure from 0.00 to 0.15 MPa when incubated in batch, determining an affinity constant (K_m_) for CH_4_ at least higher than 0.075 MPa (1.1 mM) (11). The *K_m_* for CH_4_ of ANME-2 present in Gulf of Cadiz sediment is about 37 mM (9), which is equivalent to 3 MPa CH_4_ partial pressure. This ANME-2 dominated sediment has its optimum pressure at the in situ pressure (S. Bhattarai, Y. Zhang, and P.N.L Lens, submitted for publication). In contrast, the CH_4_ partial pressure influenced the growth of different subtypes of ANME-2 and SRB (i.e. at 10.1 MPa only the ANME-2c and SEEP-SRB2 subtypes were enriched) from the Eckernförde Bay marine sediment incubated for 240 days in a high-pressure membrane capsule bioreactor (12).

Studying the effect of pressure on ANME and SRB will thus help understand the growth of the different ANME clades and their SRB partner. Finding ANME-SRB consortia that can grow fast at ambient pressure is of great importance for the application of AOM-SR in the desulfurization of industrial wastewater. Sulfate and other sulfur oxyanions, such as thiosulfate, sulfite or dithionite, are contaminants discharged in fresh water by industrial activities such as food processing, fermentation, coal mining, tannery and paper processing. Biological desulfurization under anaerobic conditions is a well-known biological treatment, in which these sulfur oxyanions are anaerobically reduced to sulfide (13–15). The produced sulfide precipitates with the metals, thus enabling their recovery (16). In the process of groundwater, mining or inorganic wastewater desulfurization, electron donor for sulfate reduction needs to be supplied externally. Electron donors such as ethanol, hydrogen, methanol, acetate, lactate and propionate (13) are usually supplied, but these increase the operational and investment costs (16). The use of easily accessible and low-priced electron donors such as CH_4_ is therefore appealing for field-scale applications (17). Moreover, from a logistic, economical and safety view point, bioreactors operating at ambient conditions are preferred over those operated at high pressures.

Coastal marine sediment from Lake Grevelingen (the Netherlands) hosts both ANME and SRB (18). Among the ANME types, ANME-3 is predominant, which makes this sediment a beneficial inoculum to investigate the effects of pressure on ANME-3. ANME-3 is often found in cold seep areas and mud volcanoes with high CH_4_ partial pressures and relatively low temperatures (10, 19, 20). Therefore, the shallow marine sediment from Lake Grevelingen was incubated at different pressures (0.1, 0.45, 10, 20, and 40 MPa) to study the influence of pressure on the AOM-SR activity, but also on the methanogenic activity and the potential formation of carbon, e.g. acetate or methanethiol, (21) and sulfur, e.g. elemental sulfur or polysulfides (22), intermediates. Moreover, phylogenetic analysis and visualization of microorganisms by fluorescence in-situ hybridization (FISH) were used to study the activity and the shifts in cell morphology, community composition and aggregation upon incubation of marine Lake Grevelingen sediment at different pressures in batch for 77 days.

## Results

### Conversion rates of sulfur compounds

The highest sulfide production rates of the coastal marine Lake Grevelingen sediment was in the incubations at the *in situ* pressure (0.45 MPa) and 10 MPa: 270 and 258 μmol g_VSS_^-1^ d^-1^, respectively (Figure 1a). The sulfide production rate at 40 MPa was 109 μmol g_VSS_^-1^ d^-1^, comparable to the rate with no CH_4_ in the headspace, 99 μmol g_VSS_^-1^ d^-1^ (Figure 1a). Similarly, high SR rates were recorded for the incubations at 0.45 MPa and 10 MPa (Figure 1b): 297 and 278 μmol g_VSS_^−1^ d^−1^, respectively. In contrast, the SR rate at 0.1 MPa was 257 μmol g_VSS_^-1^ d^-1^, while the sulfide production was only 157 μmol g_VSS_^-1^ d^-1^ (Figures 1b and 1a).

**Figure 1.**
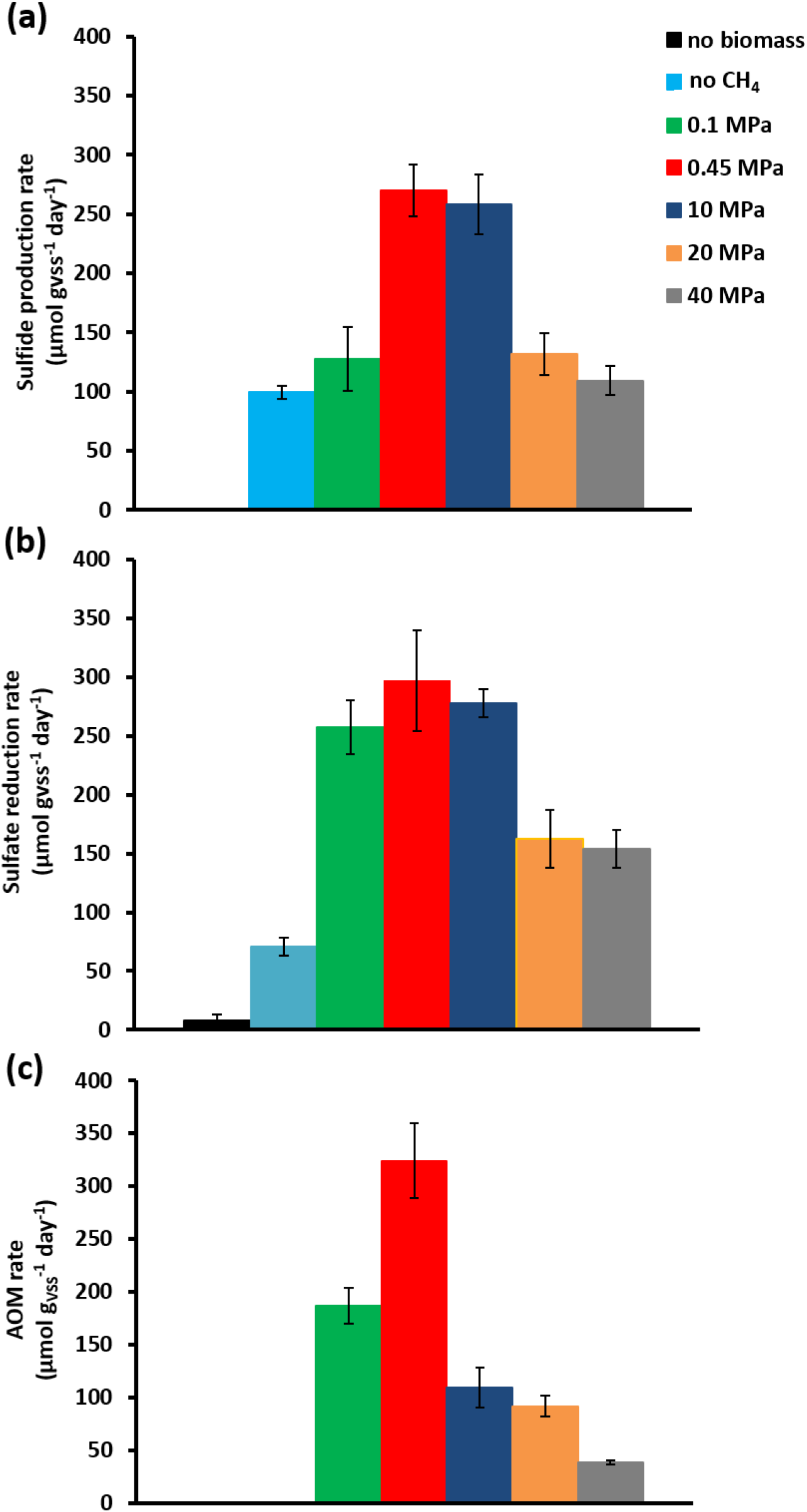
(a) Sulfide production rate, (b) SR rate and AOM rate for incubations at different pressure and controls without CH_4_, but with N_2_ in the headspace and without biomass. Error bars indicate the standard deviation (n=3).

**Table 1.**
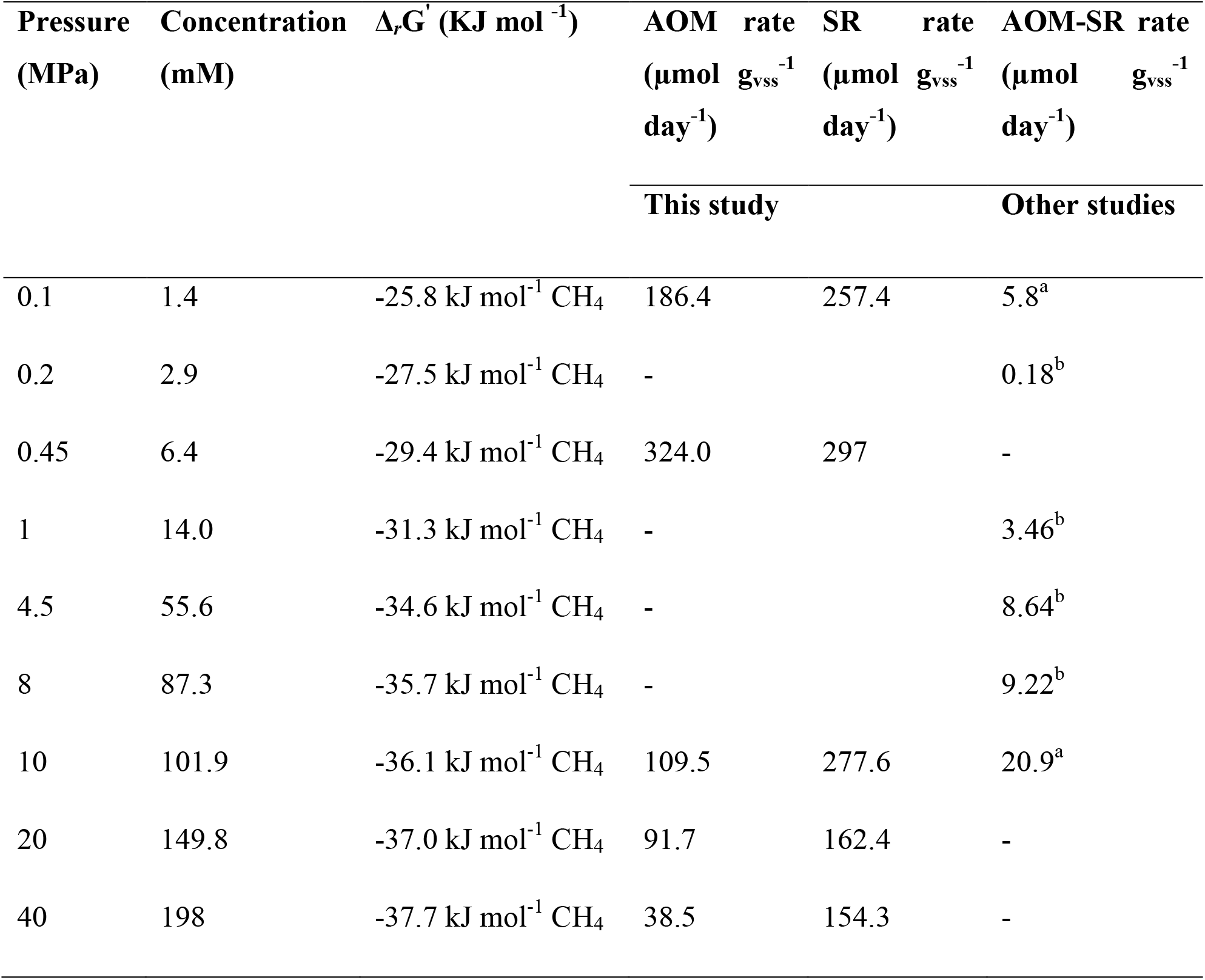
Gibbs free energy of AOM coupled to SR (Δ_*r*_G’) at different CH_4_ total pressures and assuming the following *in vitro* conditions: temperature 15°C, pH 7.0, HCO_3_ 30 mM, SO_4_^2-^ 10 mM and HS 0.01 mM. The maximum dissolved CH_4_ concentration at a salinity of 32%o and 15°C at different CH_4_ partial pressure was determined by the Duan model (52). AOM and SR rates determined in this (marine Lake Grevelingen sediment) and other studies. ^a^ = data from Timmers et al. (12): AOM-SR rate of Eckernförde sediment based on ^13^C-carbonate species production and ^b^ = data from Zhang et al. (9): AOM-SR activity of Capt Aryutinov Mud Volcano sediment based on sulfide production.

Sulfide was produced in almost all the incubations (Figure 2), with the exception of the incubation without biomass (Figure 2g). The sulfate concentration profiles varied with initial incubation pressure: at 0.45 MPa sulfate was reduced to sulfide in a 1:1 ratio (Figure 2b), whereas sulfate was not reduced anymore after 40 days of incubation at 40 MPa (Figure 2e). At 0. 45 MPa, 0.98 mmol of sulfate was consumed and exactly 0.98 mmol of total dissolved sulfide was produced, closing the sulfur balance. In the incubation at 0.1 MPa, 0.37 mmol of elemental sulfur was produced along with 0.54 mmol of sulfide (Figure 2a). In the other incubations at different pressures, hardly any elemental sulfur was formed (Figures 2c-2g). Instead, long chain polysulfides were formed along the incubation depending on the pressure (Figure 3), but in small amounts (≤ 2 μmol per vessel): 2 μmol S_6_^2-^ per vessel was determined at 0.45 MPa CH_4_ pressure (Figure 3b) and 1.2 and 1.4 μmol S_6_^2-^ per vessel at 10 MPa and 20 MPa, respectively (Figures 3c and 3d).

**Figure 2.**
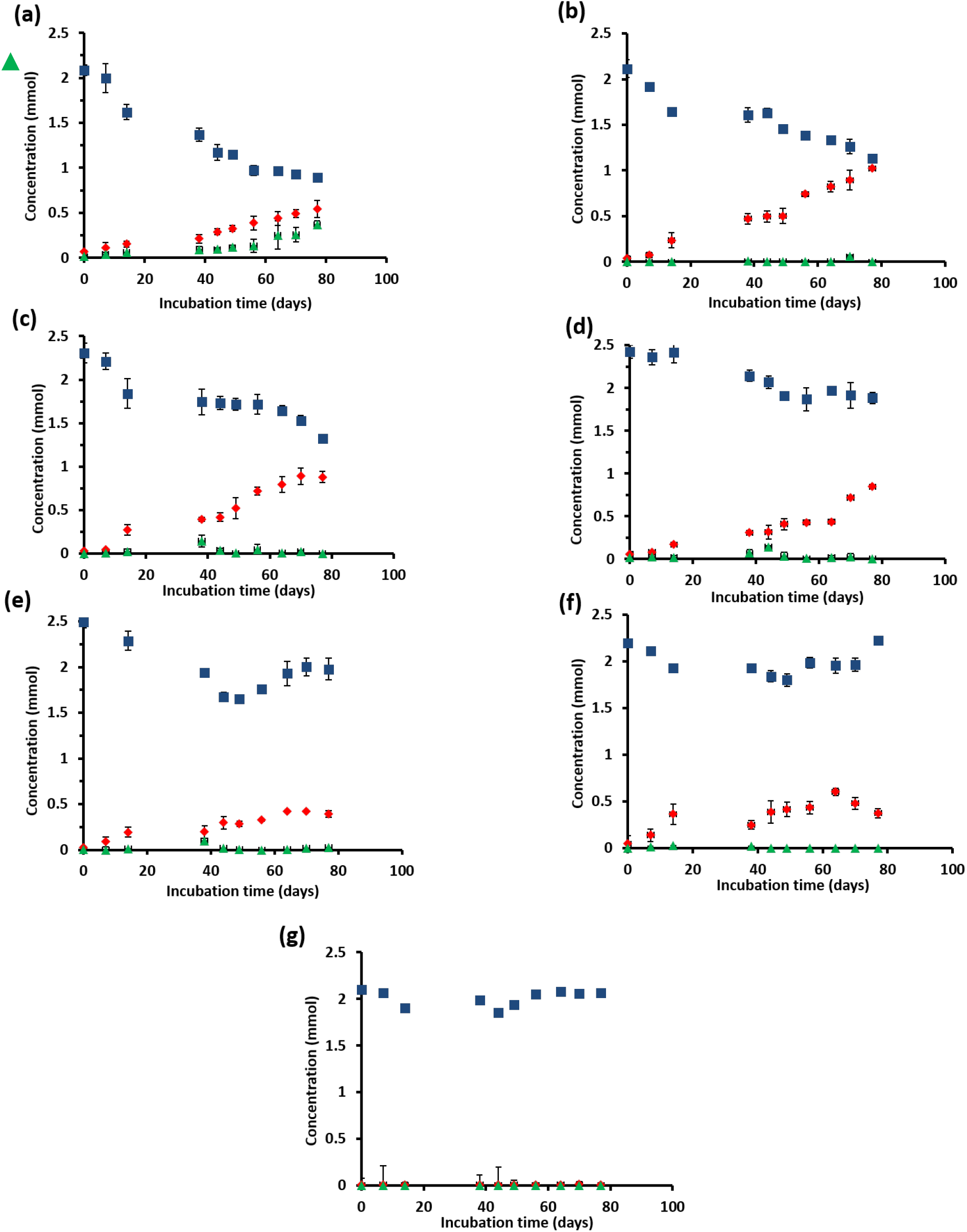
Concentration profiles of total dissolved sulfide 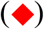, sulfate 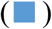 and elemental sulfur 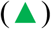 for the incubation at (a) 0.1MPa, (b) 0.45 MPa, (c) 10 MPa, (d) 20 MPa, (e) 40 MPa, (f) without CH_4_, and (g) without biomass. Error bars indicate the standard deviation (n=3).

**Figure 3.**
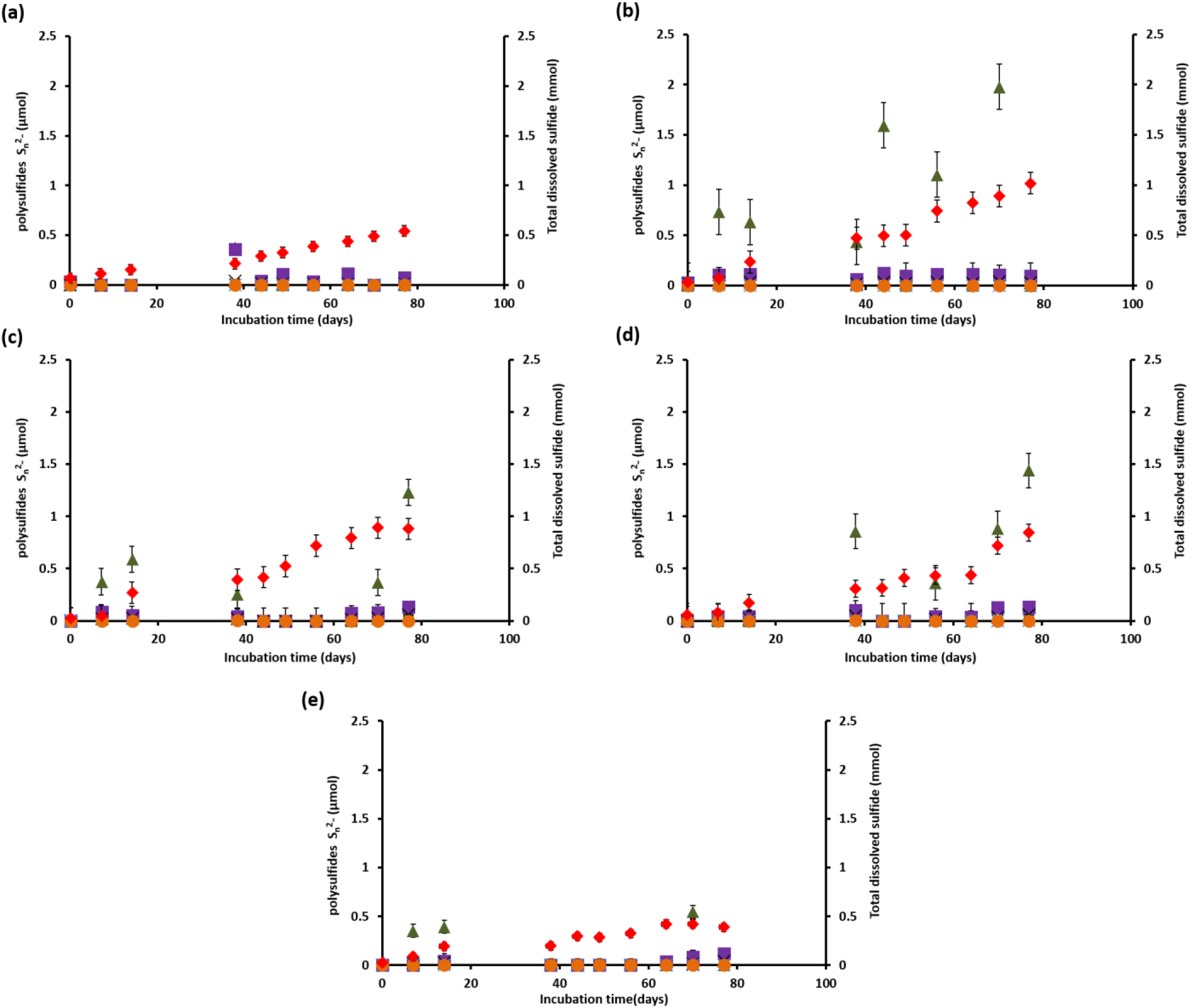
Total dissolved sulfide 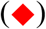 and polysulfides concentration, namely S_2_^2-^ 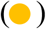, S_3_^2^’ 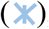, S_4_^2-^ 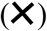, S_5_^2-^ 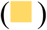. S_6_^2-^ 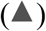, during the incubation of Grevelingen sediment at (a) 0.45 MPa, (b) 0.1MPa, (c) 10 MPa, (d) 20 MPa, and (e) 40 MPa. Error bars indicate the standard deviation (n=3).

### AOM rates

The AOM rates were calculated from the DIC produced from ^13^CH_4_, from which the *K_m_* for CH_4_ of the marine Lake Grevelingen sediment was determined to be around 1.7 mM. The DIC production rates followed a similar trend as the sulfide production rates: the highest rate was found at 0.45 MPa and the lowest rate at 40 MPa: 320 and 38 μmol g_VSS_^-1^ d^-1^, respectively (Figure 1c). In the incubation at 0.45 MPa (*in situ* pressure), the total DIC produced from CH_4_ was similar to the sulfide produced: ∼0.9 mmol per vessel (Supporting information, Figure S1b and Figure 2b). However, sulfide was produced from the start for all the incubations, while the total DIC from CH_4_ was mainly produced only after 40 days of incubation (Figure 2 and Supporting information, Figure S1). Similar trends were found for all the other incubations at different pressure, except for the vessel without CH_4_, where only sulfide production (0.3 mmol/ vessel) was recorded.

### Methanogenesis

CH_4_ was produced in all the incubations, with the exception of the batches without biomass (Supporting information, Figure S2). The highest amount of CH_4_ formed was recorded in the vessel at 0.1 MPa (Supporting Information, Figure S2b). The highest methanogenic rate was determined in the control vessel without CH_4_ (N_2_ in the headspace) and at 0.1 MPa: 44 and 31 μmol g_VSS_^-1^ d^-1^, respectively, while it was below 5 μmol g_VSS_^-1^ d^-1^ in all the other batch incubations (Supporting information, Figure S2a). Assuming that all the total ^12^C-DIC was produced from the oxidation of other carbon sources than CH_4_, its production rate was low in almost all the incubations: lower than 3 μmol g_VSS_^-1^ d^-1^, except for the incubation without CH_4_ (64 μmol g_VSS_^-1^ d^-1^, data not shown).

### Community shifts as a function of incubation pressure: total cell numbers

The total bacterial and archaeal cellular numbers were accessed from Q-PCR data performed on samples after 77 days of incubation (Figure 4). The highest increase in active cells, from 6 to 8×10^7^ cells ml^-1^, was found in the incubation at the *in situ* pressure of 0.45 MPa. In the incubation at 40 MPa, the amount of active total bacteria and archaea cells decreased from 6.5 to 5.8×10^7^ cells ml^-1^ (Figure 4a). Based on Q-PCR results, archaea grew in all the incubations, while copy numbers of bacteria decreased in the incubation without CH_4_ and at 20 MPa. The total number of archaea increased the most in the incubation at 20 MPa (Figure 4b).

**Figure 4.**
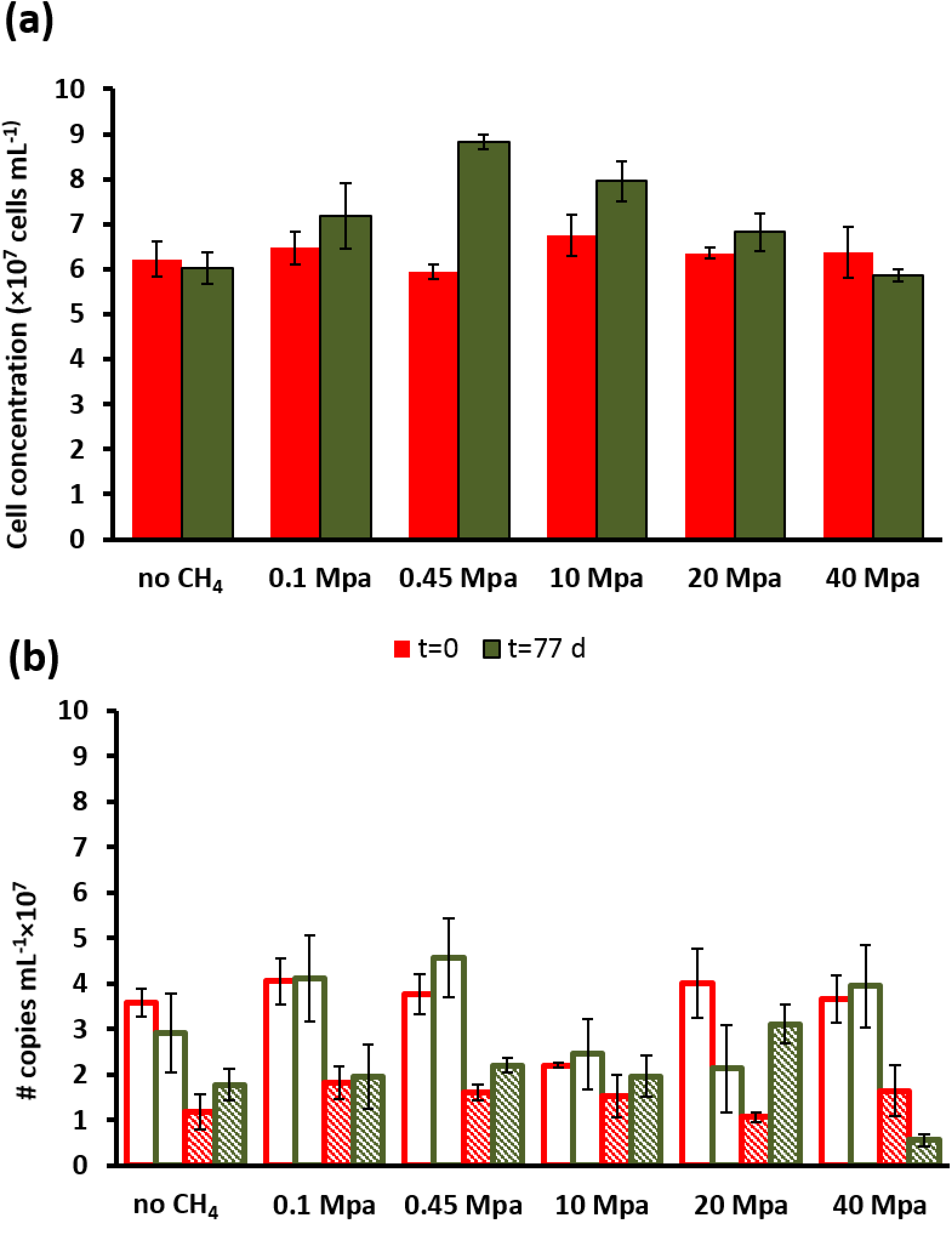
(a) Total number of active cells and (b) number of copies of archaea and bacteria from Q-PCR analysis per ml of wet sediment in each pressurized vessel at the start (t=0 days) and at the end of the incubation (t=77 days). Error bars indicate the standard deviation (n=3).

Based on the 16s rRNA gene analysis, both archaeal and bacterial communities had shifted along the 77 days incubation (Figure 5). The most abundant OTUs (operational taxonomic unit) with archaeal signature are shown in Figure 5a. Specifically, the abundance of ANME-3 among all the archaea increased the most in incubations at 0.45 and 0.1 MPa, i.e. respectively three and two times more than at the start of the incubation (Figure 5a). ANME-2a/b reads increased the most at 20 MPa: 27 times more than at the start of the incubation (Figure 5a). Sequences of methanogens, specifically belonging to the *Methanomicrobiales,* were more abundant after the incubation at 0.1 MPa, rather than at higher partial pressures, where *Thaumarchaeota* and *Woesearchaeaota* were more abundant in incubations at 10, 20 and even 40 MPa (Figure 5a).

**Figure 5.**
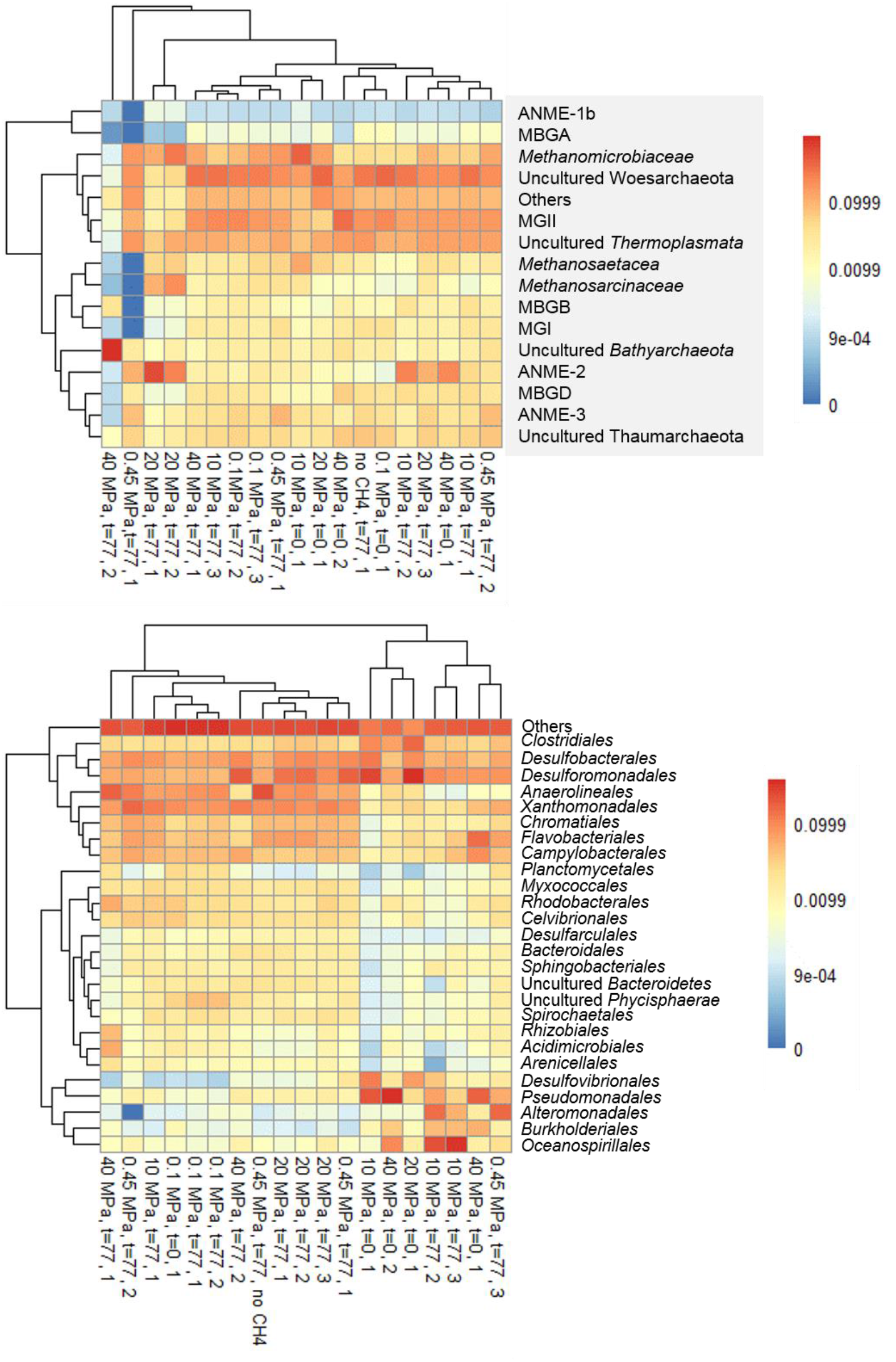
Heat map of top most abundant 16s rRNA sequences at the beginning (t=0) and at the end of the incubations (t=77 days) of the marine Lake Grevelingen sediment at different CH_4_ pressures and control without CH_4_ in the headspace showing the phylogenetic affiliation up to family level as derived by high throughput sequencing of (a) archaea and (b) bacteria.

The bacterial communities were very diverse in all the incubations, the ones with the highest percentage are shown in Figure 5b. The absolute abundance of the *Desulfobulbaceae* (DBB) as calculated from their 16s rRNA gene according to Q-PCR and Miseq results increased or remained similar at the lower pressure incubations (0.1 and 0.45 MPa), but the percentage of DBB in the total bacterial community decreased at more elevated pressures (10, 20 and 40 MPa). Differently, the absolute abundance of *Desulfobacteraceae,* as DSS, increased in all the incubations at different pressures, with the highest percentage of reads retrieved in the incubation at 20 MPa (Figure 5b). The percentage of OTUs as assigned to *Desulfovibrio, Desulfuromonas, Halomonas* and *Sulfurovum* genes decreased in all the batch incubations (Figure 5b).

### Community shifts as a function of incubation pressure: FISH analysis

ANME-3 and DBB were visualized in all the batch incubations (Figures 6a-6c). At the start of the incubation (t = 0 days), ANME-3 cells were preferentially visualized in aggregates with other cells (data not shown). FISH images after 77 days of incubation showed variations in the aggregate morphology depending on the incubation pressure (Figure 6). At 0.1 and 0.45 MPa, ANME-3 was more abundant than at the start, while the DBB cells were not found concomitant to the ANME-3 cells (Figure 6a) and, even if present, the ANME-3 cells outnumbered the DBB cells (Figure 6b). In the 10 MPa incubation, ANME-3 was visualized more scattered and not in clusters as at the lower pressures, whereas DBB cells were even more rarely pictured (Data not shown). At 20 MPa, ANME-3 and DBB cells were rare, however, the stained cells formed tight ANME-3/DBB aggregates (Figure 6c). At 40 MPa, ANME-3 and DBB were the least abundant and scattered and no aggregates could be found (data not shown).

**Figure 6.**
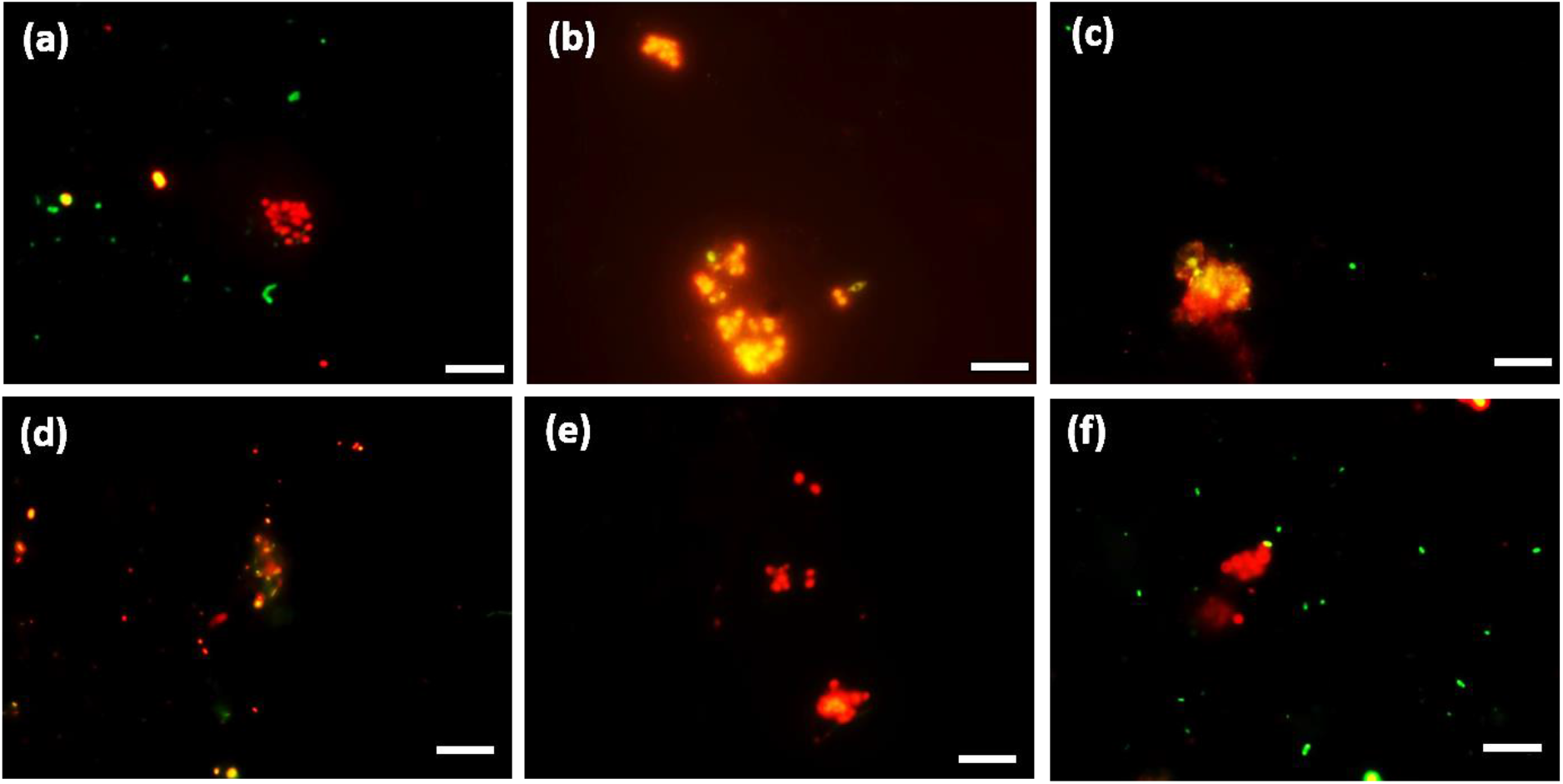
FISH images (a-c) from CY3-labeled ANME-3 in red color, CY5-labeled *Desulfobulbus* (DBB) in green after 77 days of incubation at (a) 0.1 MPa, (b) 0.45 MPa and (c) 20 MPa total CH_4_ pressure. FISH images (d-f) from CY3-labeled ANME-2 in red color, CY5-labeled *Desulforsarcina/Desulfococcus* group (DSS) in green after 77 days of incubation at (a) 0.45 MPa, (b) 20 MPa and (c) 40 MPa total CH_4_ pressure. White scale bar representing 10 μm.

Differently than ANME-3, more ANME-2 cells were visualized in the 77 days incubations at higher (10, 20 and 40 MPa) than at lower (0.1 and 0.45 MPa) incubation pressures (Figures 6d-6f). DSS, the most common SRB bacterial partner of ANME-2, were most abundant at 0.1 MPa. At lower pressure they were mainly visualized together with ANME-2 (Figure 6d). At 20 MPa only clusters of ANME-2 cells were visualized without DSS (Figure 6e).

## Discussion

### Pressure effect on AOM in marine Lake Grevelingen sediment

This study showed that AOM and SR processes in Lake Grevelingen sediment depend on the CH_4_ total pressure. According to Eq. 1, the reaction rate is expected to be stimulated by the elevated CH_4_ partial pressure when the other parameters remain the same (Table 1). This expectation has been commonly accepted and has been shown in communities dominated by ANME-1, i.e. hydrocarbon seep in the Monterey canyon sediment (23) and ANME-2, i.e. Eckernförde Bay (24, 25) and Gulf of Cadiz sediment (S. Bhattarai, Y. Zhang, and P.N.L. Lens, submitted for publication). In contrast, the AOM-SR process by the ANME-3 dominated marine

Lake Grevelingen sediment has an optimal pressure at 0.45 MPa among all tested conditions (Figure 1). This is in accordance with their natural habitat, i.e. the *in situ* pressure of marine Lake Grevelingen is 0.45 MPa, but contrasts the theoretical thermodynamic calculation (Table 1), that predicts a higher CH_4_ solubility and thus a higher activity based on reported *Km* values, i.e. 37 mM as calculated from an ANME-2 predominant enrichment originated from the Gulf of Cadiz (9). The calculated *K_m_* value on CH_4_ based on our ANME-3 dominated inoculum is much lower than previously reported: around 1.7 mM. Thus, the ANME cells from Grevelingen marine sediment have a higher affinity for CH_4_ than the ANME-2 from the Gulf of Cadiz, explaining why a higher pressure, i.e. higher dissolved CH_4_ concentration, did not result in a higher AOM rate (Figu 1c). As a matter of fact, higher pressure resulted in a lower number of ANME-3 cells and ANME-3 proliferated only at lower pressures incubation (0.1 and 0.45 MPa). This indicates ANME-3 cells of marine Lake Grevelingen are non-piezophilic, which are easily damaged upon pressure elevation and require extra energy to cope with the damage (26). ANME-3 are found in cold seep areas and mud volcanoes with high CH_4_ partial pressures (∼10 MPa) and relatively low temperatures (10, 19, 20). However, Lake Grevelingen is a shallow sediment with high abundance of ANME-3 (18) and likely contains different subtypes than the ones found in deep sea sediments, which cannot cope with a high pressure.

### Pressure effect on ANME types

The ANME-3 type is usually visualized in association with DBB as sulfate reducing partner (10, 19). FISH analysis showed that the DBB cells were not as high in number as the ANME-3 cells in any of the incubations (Figures 6a-6c), but they increased the most at the 0.1 MPa incubation (data not shown). In a recent study describing the microbial ecology of Lake Grevelingen sediment (incubation pressure = 0.1 MPa), the two species (ANME-3 and DBB) could not be visualized together and the DBB cells were much less abundant than ANME-3 (18), similarly to this study (Figure 6a). At 0.1 and 0.45 MPa, ANME-3 cells were visualized in aggregates mainly detached from DBB cells (Figures 6a and 6b). ANME-3 cells have been visualized without bacterial partner before (20, 27), suggesting that this ANME type is supporting a metabolism independent of an obligatory bacterial association. In contrast, as ANME-3 and DBB decreased in number at higher pressures, most of the ANME-3 and DBB visualized at 20 MPa were forming small ANME-3/DBB clusters, suggesting that they have mutual benefits at this pressure (Figure 6c).

Sequences of ANME-2 were also found by Miseq analysis (Figure 5a) and visualized by FISH (Figures 6d-6f) in all incubations. ANME-2a/b cells were higher in number in the incubation at higher pressures (10 and 20 MPa). Also many DSS were found in all the batch incubations and, as for ANME-2, they were more abundant at higher pressures (10 and 20 MPa). ANME-2 and DSS were mainly visualized in aggregates (Figure 6d), especially at lower pressures (0.1 and 0. 45 MPa). The cooperative interaction between the ANME-2 and DSS is still under debate: Milucka et al. (22) stated that a synthrophic partner might not be required for ANME-2 and that they can be decoupled by using external electron acceptors (28), whereas recent studies have shown direct electron transfer between the two (ANME-2 and DSS) partners (29, 30). Besides, the DSS might have proliferated by growth on organic carbon compounds released by damaged or killed piezosensitive microorganisms.

### Effect of pressure on sulfur cycle in marine Lake Grevelingen sediment

Figures 2 and 3 shows that the total pressure steers the sulfur cycling in the marine Lake Grevelingen sediment community. At 0.1 MPa CH_4_ pressure, the reduced sulfate was converted to both sulfide and zero-valent sulfur (Figure 2a). At 0.45 MPa (the incubation with the highest AOM-SR activity), the sulfur balance was closed by solely the sulfide production (Figure 2b). The production of elemental sulfur was repressed at elevated CH_4_ pressures (Figure 2b-2e). Elemental sulfur has been considered as intermediate in the AOM-SR process, which is consumed by ANME to generate energy (22). Milucka et al. (22) showed that ANME-2 cells could stand along without the metabolic support of the bacterial partner, assuming that CH_4_ was oxidized to bicarbonate and sulfate was reduced to disulfide (S_2_^2-^) through zero-valent sulfur as an intracellular intermediate. The amount of disulfide or other polysulfides formed during the incubations (Figure 3) was very low, in most cases below the detection limit (0.1 μmol). The formation of these intermediate sulfur compounds in the ANME process needs to be further elucidated using e.g. isotopic labeled sulfate (^35^S) and nanometre scale secondary ion mass spectrometry (NanoSIMS) analysis.

A shift from sulfate reducers (e.g. *Desulfobacterales*) to sulfur reducers (e.g. *Desulforomonadales*) was observed in the bacterial community from low to high CH_4_ partial pressure (Figure 5b). Sulfur reducing bacteria, e.g. *Desulfovibrio* or *Desulforomonas*, are more abundant at high CH_4_ partial pressure (10, 20, 40 MPa), whereas sulfate reducing DBB are more abundant in the incubations at lower CH_4_ total pressure (Figure 5b), where they were present in ANME-DBB aggregates and had the highest AOM-SR rates (Figure 1).

### In vitro demonstration of SR-AOM supported ecosystem in Lake Grevelingen

This study showed that CH_4_ and sulfate were an effective energy source supporting SR-AOM in the microbial ecosystem from the marine Lake Grevelingen sediment (Figure 1). Apparent *in vitro* biomass growth was observed, especially at 0.45 MPa which is the *in situ* pressure, with

CH_4_ and sulfate supplied as the sole energy sources (Figure 1). At incubation conditions similar to *in situ* conditions (p = 0.45 MPa, T = 15ºC, pH = 7), the AOM and SR rates reached approximately 0.3 mmol g_VSS_^-1^ d^-1^. These rates are comparable or even higher than the *in vitro* AOM rates of ANME-1 or ANME-2 dominated biomass, e.g. the rate obtained after the enrichment of Eckernförde Bay sediment dominated by ANME-2 type cells for more than 800 days in a continuous membrane bioreactor (31). Moreover, the AOM-SR rate measured in this study at 0.45 MPa is even higher than the AOM rate coupled to denitrification, which is thermodynamically much more favorable (ΔG^0’^ = −924 kJ mol^-1^ CH_4_) (32) than AOM-SR (ΔG^0’^ = −16.6 kJ mol^-1^ CH_4_, Eq. 1).

It should be noted that even after two months incubation, the abundance of the responsible microorganisms, i.e. all detected types of ANME and SRB cells, is quite low: 17.8 ×10^5^ and 11.4 ×10^5^ number of copies per mL of wet sediment of ANME-3 and ANME-2, respectively, in the total community (data not shown). The ANME-3 cells present in the marine Lake Grevelingen possess high specific AOM-SR rates and thus, can be of great potential to be applied in the industry after enrichment. The SR rate with CH__4__ as electron donor should be around 100 mmol g_VSS_^-1^ d^-1^ to be competitive with the SR rates achieved with other electron donors, such as hydrogen or ethanol (31, 33), which is still much higher than what was obtained in this study.

Methanogenic activity in marine Lake Grevelingen sediment was previously described by Egger et al. (34) and confirmed in this study at low pressure (0.1 MPa) or when no CH_4_ was added in the incubation (Supporting Information, Figure S2). At 0.1 MPa, the CH_4_ production rate was 31 μmol g_VSS_^-1^ d^-1^ and the AOM rate was 186 μmol g_VSS_^-1^ d^-1^. Trace CH_4_ oxidation occurs during methanogenesis and the archaea involved compete with SRB for acetate (12, 24). Thus, the determined AOM at 0.1 MPa cannot account for the net AOM-SR, as it is partly due to the methanogenic activity.

At high pressures (0.45, 10 MPa), AOM-SR was preferred (Figure 1) over methanogenesis (Supporting Information, Figure S2). Methanogenesis becomes less thermodynamically favorable at high pressures, less free energy (12 kJ mol^-1^ less) is released upon changing the incubation pressure from 0.1 to 10 MPa (24). Timmers et al. (12) found that at 10 MPa net AOM-SR occurred, while at 0.1 MPa methanogenesis and trace CH_4_ oxidation dominated. In this study, the optimal AOM-SR was 0.45 MPa: the SR activity decreased at pressures higher than 10 MPa, while AOM activity already decreased at pressures higher than 0.45 MPa (Figures 1b and 1c).

## Conclusions

This is the first study showing that the active ANME from the shallow marine Lake Grevelingen sediment preferred lower (0.1 and 0.1 MPa) over elevated (10, 20, 40 MPa) pressures, in contrast to previous studies that show strong positive correlations between the growth of ANME-1/2 and the CH_4_ pressure. Pressure steered the abundance and structure of the different types of ANME and SRB. The ANME-3 type was predominantly enriched in incubations at low pressures, whereas high pressures enhanced ANME-2 proliferation. Similarly, a shift from sulfate reducers to sulfur reducers was observed in the bacterial community from low (0.1 and 0.45 MPa) to high (10, 20, 40 MPa) CH_4_ partial pressure. This research highlights that ANME-3 from marine Lake Grevelingen can be enriched at rather low CH_4_ partial pressures, which is important to further understand their metabolism and physiology.

## Materials and Methods

### Site description and sampling procedure

The sediment was obtained from the Scharendijke Basin in the marine Lake Grevelingen (water depth of 45; position 51° 44.541’ N; 3° 50.969’ E), which is a former estuary in the southwestern part of the Netherlands. The sampling site characteristics, biochemical processes and the microbial community composition have been described previously (18, 34-36). Coring was done in November 2013 on the vessel R/V Luctor by the Royal Netherlands Institute for Sea Research (Yerseke, the Netherlands). The sampling procedure has been described in Bhattarai et al. (18), the sediment was kept at 4 °C in the dark in serum bottles with a headspace of CH_4_ until use.

### Experimental design

The effect of the pressure on the CH_4_ oxidation, SR and CH_4_ production rate of the marine Lake Grevelingen sediment was assessed with 0.07 (± 0.01) g volatile suspended solids (g_VSS_) in 200 ml pressure vessels incubated in triplicates at 0.1 MPa, 0.45 MPa (mimicking the in situ conditions), 10 MPa, 20 MPa and 40 MPa. The vessels were flushed and pressurized with 100 % CH_4_, from which about 20% was ^13^C-labeled CH_4_ (^13^CH_4_). The marine Lake Grevelingen sediment used as inoculum was incubated with artificial saline mineral medium with sulfate (10 mM) at 15°C for 77 days. Two different control incubations were prepared in triplicates at 0.45 MPa: without biomass and without CH_4_, but with nitrogen in the headspace.

Slurry samples were taken every week for chemical analysis. Approximately 1 mL sample was taken by attaching a connector and a vacuum tube to the exit port while gently opening the tap. Weight and pressure were measured in the vacuum tube before and after sampling. Pressure in each vessel was restored by adding fresh basal medium using a high performance liquid chromatography (HPLC) pump (SSI, USA).

### Chemical analysis

The gas composition was measured on a gas chromatograph-mass spectrometer (GC-MS Agilent 7890A-5975C). The GC-MS system was composed of a Trace GC equipped with a GC-GasPro column (30 m by 0.32 mm; J & W Scientific, Folsom, CA) and an Ion-Trap MS. Helium was the carrier gas at a flow rate of 1.7 ml min^-1^. The column temperature was 30ºC. The fractions of CH_4_ and CO_2_ in the headspace were derived from the peak areas in the gas chromatograph, while the fractions of^13^ CH_4_, ^12^CH_4_, ^12^CO_2_ and ^13^CO_2_ were derived from the mass spectrum as done by (37).

Total dissolved sulfide was measured by using the methylene blue method (Hach Lange method 8131) and a DR5000 Spectrophotometer (Hach Lange GMBH, Düsseldorf, Germany). Samples for sulfate and thiosulfate analysis were first diluted in a solution of zinc acetate (5 g/L) and centrifuged at 13,200g for 3 min to remove insoluble zinc sulfide, and filtrated through 0.45 μm membrane filters. Sulfate and thiosulfate concentrations were then determined by ion chromatography (Metrohm 732 IC Detector) with a METROSEP A SUPP 5 – 250 column. The pH was checked by means of pH paper.

Polysulfides were methylated using the protocol by Kamyshny et al. (38) and analyzed by reversed-phase HPLC. Elemental sulfur from the slurry sample was extracted using methanol following the method described by Kamyshny et al. (39), but modified for small volumes. Dimethylpolysulfanes and extracted elemental sulfur were analyzed by an HPLC (HPLC 1200 Series, Agilent Technologies, USA) with diode array and multiple wavelength detector. A mixture of 90% MeOH and 10% water was used as eluent. A reversed phase C-18 column (Hypersil ODS, 125 × 4.0 mm, 5 μm, Agilent Technologies, USA) was used for separation.

Concentrations of dimethylpolysulfanes from Me_2_S_3_ to Me_2_S_7_ were calculated from calibration curves of polysulfides standards prepared following the protocol of Milucka et al. (22). UV detector response to Me_2_S_8_ was calculated by the algorithm discussed in Kamyshny et al. (40).

The VSS was estimated at the beginning of the experiment on the basis of the difference between the dry weight total suspended solids and the ash weight of the sediment according to the procedure outlined in Standard Method (41).

### Rate calculations

Both AOM and SR rates were expressed as μmol of sulfide or dissolved inorganic carbon (DIC) production per gram of VSS per day (μmol g_VSS_^-1^ d^-1^). For the AOM rate calculation, the total production of ^13^C-carbonate species (^13^ C-DIC), i.e. ^13^CO_2_ in both liquid and gas phases, H ^13^CO_3_^-^ and ^13^CO_3_^2-^ in liquid phase, were first calculated. Considering that only 20% of CH_4_ was ^13^CH_4_, the total ^13^C-DIC was divided by the fractional abundance of ^13^C in the CH_4_ measured and used for each batch to determine the total amount of DIC produced from CH_4_ oxidation (42). For methanogenesis and for the formation of carbonate species from other carbon sources than CH_4_, ^12^CH_4_ and H^12^CO_3_^-^ were taken respectively, and divided by the ^12^C fractional abundance. A line was plotted over the period where the decrease or increase of the different compounds (^12^CH_4_, ^13^CH_4_, H^12^CO_3_^-^, H^13^CO_3_^-^, total dissolved sulfide and sulfate) was linear (at least four consecutive points) to estimate the rates (24), which were divided by the biomass content in the vessels (0.07 ± 0.01 g_VSS_ in each vessel).

### DNA extraction

DNA was extracted by using a FastDNA^®^ SPIN Kit for soil (MP Biomedicals, Solon, OH, USA) by following the manufacturer’s protocol. Approximately 0.5 g of the sediment was used for DNA extraction from the initial inoculum and ∼0.5 ml of liquid obtained by washing the polyurethane foam packing with nuclease free water was used for extracting DNA from the enriched slurry. The extracted DNA was quantified and quality was checked as described by Bhattarai et al. (18).

### PCR amplification for 16S rRNA genes and Illumina Miseq data processing

The DNA was amplified using bar coded archaea specific primer pair Arc516F and reverse Arc855R. The PCR reaction mixture was prepared as described by Bhattarai et al. (18), however, the PCR amplification was performed using a touch-down temperature program. PCR conditions consisted of a pre-denaturation step of 5 min at 95°C, followed by 10 touch-down cycles of 95°C for 30 sec, annealing at 68°C for 30 sec with a decrement per cycle to reach the optimized annealing temperature of 63°C and extension at 72°C. This was followed by 25 cycles of denaturation at 95°C for 30 sec and 30 sec of annealing and extension at 72°C. The final elongation step was extended for 10 min.

The primer pairs used for bacteria were forward bac520F 5’-3’ AYT GGG YDT AAA GNG and reverse Bac802R 5’-3’ TAC NNG GGT ATC TAA TCC (43). The following program was used: initial denaturation step at 94°C for 5 min, followed by denaturation at 94°C for 40 sec, annealing at 42 °C for 55 sec and elongation at 72°C for 40 sec (30 cycles). The final elongation step was extended to 10 min. 5 μl of the amplicons were visualized by standard agarose gel electrophoresis (1% agarose gel, a running voltage of 120 V for 30 min, stained by gel red) and documented using a UV transilluminator with Gel Doc XR System (Bio-Rad, USA).

After checking the correct band size, 150 μl of PCR amplicons were loaded in 1% agarose gel and electrophoresis was performed for 120 min at 120 V. The gel bands were excited under UV light and the PCR amplicons were cleaned using E.Z.N.A.^®^ Gel Extraction Kit by following the manufacturer’s protocol (Omega Biotek, USA). The purified DNA amplicons were sequenced by an Illumina HiSeq 2000 (Illumina, San Diego, USA) and analyzed according to the procedure described in Bhattarai et al. (18). A total of 40,000 (± 20,000) sequences were assigned to archaea and bacteria examining the tags assigned to the amplicons. More detailed analytical procedure has been described by Bhattarai et al. (18). These sequence data have been submitted to the NCBI GenBank database under BioProject accession number PRJNA415004 (direct link: http://www.ncbi.nlm.nih.gov/bioproject/415004).

### Quantitative real-time PCR (Q-PCR)

Archaeal and bacterial clones were used to prepare Q-PCR standard. Plasmids were isolated using the Plasmid Kit (Omega Biotek, USA). The plasmid was digested with the EcoR I enzyme. After digestion purification was done by gel extraction (Gel extraction Kit, Omega Biotek, USA). The copy number was calculated from the total mass and the nucleic acid concentration. Extracted DNA from the sediment at the start and at the end of the incubation period (11 weeks) was used for qPCR analysis to quantify archaea and bacteria. Amplifications were done in triplicates in a 7500 Real-Time PCR System (Applied Biosystem). Each reaction (20μl) contained 1× Power SYBR-Green PCR MasterMix (Applied Biosystems), 0.4μM of each primer, and 5 ng template DNA. The 16S rRNA genes of bacterial origin were amplified using the primers Bac331f (5’-TCCTACGGGAGGCAGCAGT3’) and Bac797r (5’-GGACTACCAGGGTCTAATCCTGTT-3’) (44). Cycling conditions were 95°C for 10 min; and 40 cycles at 95°C for 30 sec and 60°C for 30 sec and 72°C for 30 sec. Archaea were quantified using the primer set Arch349f (5’-GYGCASCAGKCGMGAAW-3’) and Arch806r (5’-GGACTACVSGGGTATCTAAT-3’) (45). Cycling conditions were 95°C for 10 min; and 40 cycles at 95°C for 30 sec and 50°C for 30 sec and 72°C for 30 sec. Triplicate standard curves were obtained with 10-fold serial dilutions ranged between 10^7^ and 10^−2^ copies per μl of plasmids. The efficiency of the reactions was up to 100% and the R^2^ of the standard curves were up to 0.999.

### Cell visualization and counting by FISH

At the start and at the end of the incubation period (11 weeks), 200 μL of sample from each vessel was fixed in a final 2% paraformaldehyde solution for 4 h on ice. The samples were washed twice with 1x phosphate buffer saline solution (PBS). Then, it was stored in a mixture of PBS and ethanol (EtOH), with a PBS/EtOH ratio of 1:1 at −20°C as previously described by (3). This sample was used for cell counting and FISH analysis.

100 μL of stored sample was diluted with nuclease free water, sonicated for 40 sec and then filtered on 0.2 μm membrane filters. For cell counting, 200-300 μL of 20x SYBR green solution (Takara, Japan) was added on top of the filter and incubated in the dark at room temperature for 30min. The filters were dried and mounted on a glass slide with 100 μL glycerol 10%. For FISH analysis, the filtrated sample was hybridized with the archaeal probe ARCH915 (46), the bacterial probe EUB I-III (47), with different CY3-labeled ANME probes: ANME-1 350 (3), ANME-2 538 (48), ANME-3 1249 (10) and the Cy5-labelled SRB specific probes for *Desulfosarcina / Desulfococcus* (DSS) DSS658 (3) and *Desulfobulbus* (DBB) DBB660 (49). Cells were counterstained with 4’, 6-diamidino-2-phenylindole (DAPI) (50). The hybridization of the samples and microscopic visualization of the hybridized cells were performed as described previously (51).

## Acknowledgements

We acknowledge Filip Meysman from the Royal Netherlands Institute of Sea Research (NIOZ, Yerseke, The Netherlands) for providing the Lake Grevelingen sediment. The authors are grateful for using the analytical facilities of the Centre for Chemical Microscopy (ProVIS) at the Helmholtz-Centre for Environmental Research which is supported by European regional Development Funds (EFRE and Europe funds Saxony) and the Helmholtz Association. The authors acknowledge Dr. Niculina Musat (team leader of ProVIS) for her advice and practical support during FISH analysis. This research was funded by the Erasmus Mundus Joint Doctorate Programme ETeCoS^3^ (Environmental Technologies for Contaminated Solids, Soils and Sediments) under the grant agreement FPA no. 2010-0009 and the National Natural Science Foundation of China (grant number: 41476123).

---

Figure S1. (a) CH_4_ production rates were calculated from the linear regression over at least four successive measurements in which the calculated ^12^CH_4_ increase over time was linear. (b) The CH_4_ produced was calculated from the ^12^CH_4_. Methanogenic activity and CH_4_ produced during AOM were determined for incubations at different pressures and controls without CH_4_, but with N_2_ in the headspace and without biomass. Error bars indicate the standard deviation (n=4).

Figure S2. Concentration profiles of methane oxidized 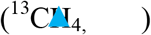 and dissolved inorganic 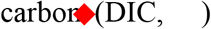 calculated from the produced ^13^CO_2_) during the incubation of marine Lake Grevelingen sediment at (a) 0.1 MPa, (b) 0.45 MPa, (c) 10 MPa, (d) 20 MPa and (e) 40 MPa.

